# Activation and modulation of the host response to DNA damage by an integrative and conjugative element

**DOI:** 10.1101/2024.10.09.617469

**Authors:** Saria McKeithen-Mead, Mary E. Anderson, Alam García-Heredia, Alan D. Grossman

## Abstract

Mobile genetic elements help drive horizontal gene transfer and bacterial evolution. Conjugative elements and temperate bacteriophages can be stably maintained in host cells. They can alter host physiology and regulatory responses and typically carry genes that are beneficial to their hosts. We found that ICE*Bs1*, an integrative and conjugative element of *Bacillus subtilis*, inhibits the host response to DNA damage (the SOS response). Activation of ICE*Bs1* before DNA damage reduced host cell lysis that was caused by SOS-mediated activation of two resident prophages. Further, activation of ICE*Bs1* itself activated the SOS response in a subpopulation of cells, and this activation was attenuated by the functions of the ICE*Bs1* genes *ydcT* and *yddA* (now *ramT* and *ramA*, for RecA modulator). Double mutant analyses indicated that RamA functions to inhibit and RamT functions to both inhibit and activate the SOS response. Both RamT and RamA caused a reduction in RecA filaments, one of the early steps in activation of the SOS response. We suspect that there are several different mechanisms by which mobile genetic elements that generate ssDNA during their lifecycle inhibit the host SOS response and RecA function, as RamT and RamA differ from the known SOS inhibitors encoded by conjugative elements.

## Introduction

Horizontal gene transfer contributes to microbial evolution by allowing bacteria to acquire new genes and phenotypes through the transfer of DNA from a donor organism to a recipient. Horizontal gene transfer is often mediated by mobile genetic elements (MGEs), including bacteriophages and conjugative elements, and many bacterial genomes contain multiple MGEs. Functional or defective temperate phages are usually (but not always) found integrated in the bacterial genome. Conjugative elements include plasmids (extrachromosomal) and integrative and conjugative elements (ICEs) that reside integrated in the host genome. Both conjugative plasmids and ICEs encode conjugation machinery, a type IV secretion system, that is capable of contact-dependent transfer of element (and sometime other) DNA from donor to recipient cells to generate transconjugants. Bacteriophages and conjugative elements often reside in and co-evolve with a bacterial host.

ICE*Bs1* is an integrative and conjugative element of *Bacillus subtilis* (Burrus and Waldor, 2004; Auchtung *et al*., 2005). It is regulated by cell-cell signaling and activated by the host SOS response to DNA damage (**Fig 1A**) ((Auchtung *et al*., 2005). As with other ICEs, ICE*Bs1* excises from the chromosome and the circular ICE DNA is nicked, unwound and undergoes rolling-circle replication (Auchtung *et al*., 2007; Lee and Grossman, 2007; Lee *et al*., 2007; Lee *et al*., 2010; Thomas *et al*., 2013). Replication is important in the original host (donor) for maintenance of the element during cell growth (Lee *et al*., 2010) and in transconjugants to allow for re-repression and integration into the genome (Lee *et al*., 2007; McKeithen-Mead and Grossman, 2023). DNA unwinding and rolling-circle replication generate the linear single stranded DNA (ssDNA) that is transferred from donor to recipient during conjugation (Lee and Grossman, 2007; Lee *et al*., 2010).

**Figure 1.**
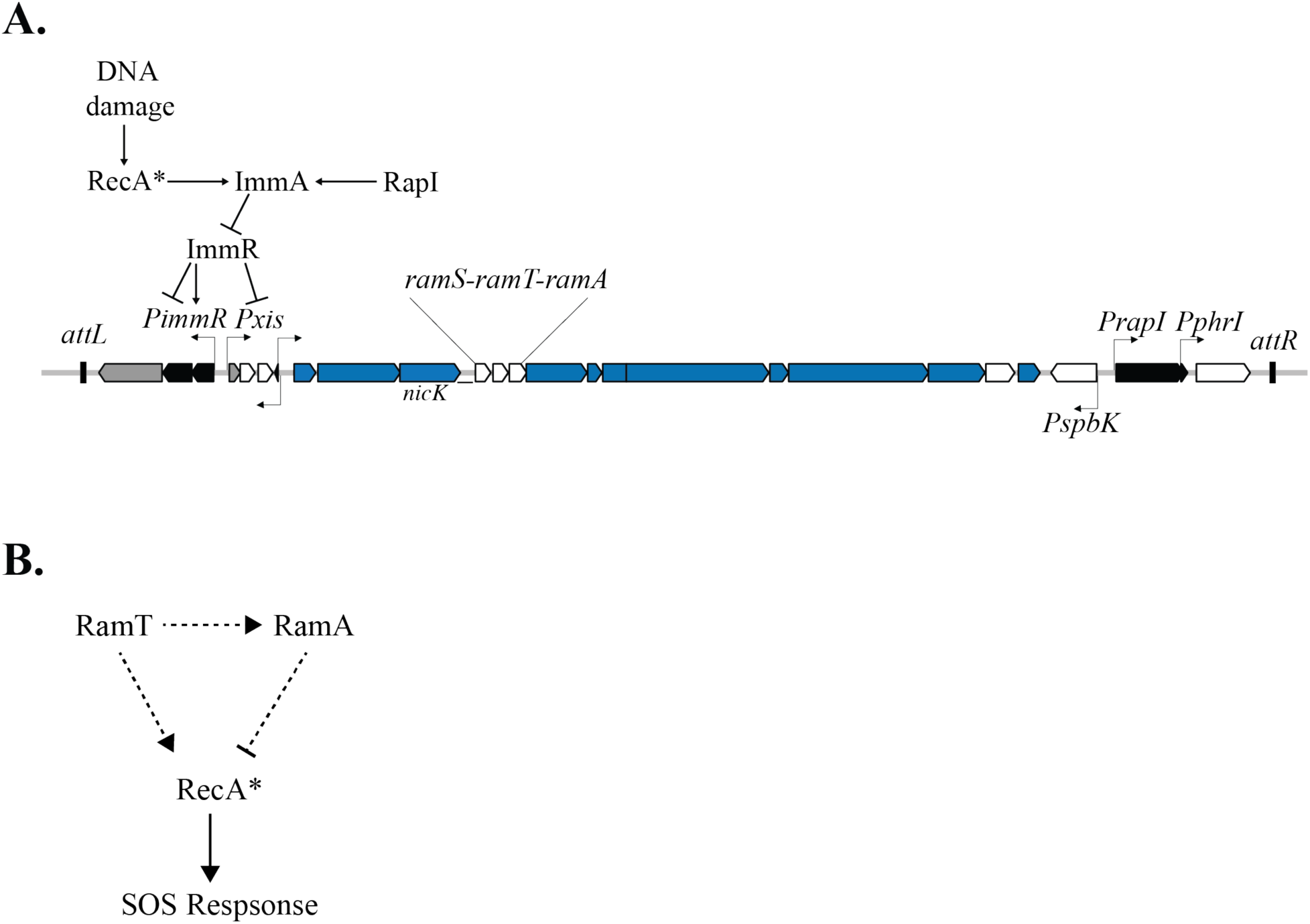
Gene map and regulation of ICE*Bs1* and model for the functions of RamT and RamA. **A.** Map of ICE*Bs1*. The junctions between the ICE (∼21 kb) and the chromosome are indicated as small rectangles at the left (*attL*) and right (*attR*) ends of the element. Genes are represented as rectangles with arrows at the end indicating the direction of transcription. Blue: genes that are required for conjugation and encode the type IV secretion system and the relaxasome; Gray: genes needed for integration (*int* at the left end) and excision (*xis*); Black: genes involved in regulation, including *immA* (protease), *immR* (repressor), and *rapI-phrI* (quorum sensing regulation); white: genes not required for conjugation, including cargo genes, genes of unknown function, and *ramS, ramT*, and *ramA* (this work; previously *ydcS, ydcT* and *yddA*, respectively). Lines with arrows above and below the gene map indicate promoters and the direction of transcription. The major promoter that drives transcription of the conjugation genes is P*xis* (Auchtung *et al*., 2005; Auchtung *et al*., 2007). A simplified cartoon of regulation is depicted above. ImmR represses transcription from P*xis*, and both activates and represses transcription from P*immR* (Auchtung *et al*., 2007). When integrated in the chromosome, P*xis* is repressed by ImmR. P*xis* expression is derepressed when ImmR is cleaved by the anti-repressor and protease ImmA (Bose *et al*., 2008), which is activated by the RecA-dependent DNA damage response or RapI. RapI is inhibited by the ICE*Bs1*-encoded peptide PhrI (not depicted) (Auchtung *et al*., 2005). **B.** Model for the functions of RamT and RamA. RamA functions to inhibit and RamT functions to both inhibit and activate the host SOS response through RecA, either directly or indirectly. RamA acts as an inhibitor of RecA. RamT both activates and inhibits RecA, and inhibition is through stimulating or enabling RamA repression. Dotted lines indicate that these interactions could be direct or more likely, indirect.

In many bacterial species, including *B. subtilis*, the presence of ssDNA is typically a signal of DNA damage and can initiate the host SOS response (Sassanfar and Roberts, 1990; Lovett Jr *et al*., 1994). Filaments of RecA form on the ssDNA, activating RecA for its roles in gene regulation and homologous recombination (Love and Yasbin, 1986; Kuzminov, 1999; Lusetti and Cox, 2002; Friedberg *et al*., 2005; Goranov *et al*., 2006). Because rolling-circle replication of ICE*Bs1* generates ssDNA that could trigger the SOS response, we postulated that the element might have genes that reduce this potential response, perhaps analogous to *psiB* of the family of F plasmids of *E. coli* (Bagdasarian *et al*., 1980; Bailone *et al*., 1988; Petrova *et al*., 2009). The *psiB* gene product reduces the SOS response in transconjugants by binding to RecA and inhibiting formation of RecA filaments on the ssDNA (Jones *et al*., 1992; Bagdasarian *et al*., 1992; Althorpe *et al*., 1999; Petrova *et al*., 2009).

We found that ICE*Bs1* inhibits the host SOS response to DNA damage in host (donor) cells. When activated, ICE*Bs1* inhibited DNA damage-induced host killing by the temperate phage SPβ and the defective temperate phage PBSX. Two genes in ICE*Bs1*, *ydcT* and *yddA* (now called *ramT* and *ramA*, for <u>R</u>ec<u>A m</u>odulator) were involved: both RamA and RamT functioned to inhibit and RamT also functioned to activate the host SOS response (**Fig 1B**). These proteins affected the accumulation of RecA filaments, an essential process in the normal cellular response to DNA damage. Additionally, activation of ICE*Bs1* itself stimulated the SOS response in a subpopulation of cells and *ramT* and *ramA* functioned to limit this activation. *ramT* and *ramA* are conserved in some ICEs and do not appear to be related to genes on other conjugative elements that are known to inhibit the SOS response. We suspect that many ICEs have genes that modulate host responses to DNA damage, perhaps in both donor and recipient cells, and that there is likely a variety of different mechanisms used.

## Results

### ICE*Bs1* reduces cell lysis caused by UV irradiation

In many species of bacteria, irradiation with ultraviolet (UV) light causes DNA damage and a global regulatory response, the SOS response (Yasbin, 1977). This response can result in cell lysis, mediated, at least in part, by de-repression of resident prophage. We monitored effects of ICE*Bs1* in *B. subtilis* on the DNA damage response by measuring cell growth and phage-mediated lysis following UV irradiation. Briefly, cells were grown in defined minimal medium to mid-exponential phase, exposed to UV light, and cell density (OD600) was monitored for several hours (Methods). Where indicated, ICE*Bs1* was activated by adding xylose to cause expression of P*xyl*-*rapI*. RapI activates ICE*Bs1* (**Fig 1A**) by causing cleavage of the element-encoded repressor ImmR by the element-encoded protease ImmA (Auchtung *et al*., 2007; Bose *et al*., 2008; Bose and Grossman, 2011).

As expected, treatment with UV light caused cell lysis. Cultures of cells containing an integrated, repressed ICE*Bs1* [ICE+ (off); SAM248] or cells without ICE*Bs1* [ICE(0); SAM591] grew normally without UV irradiation, but had a precipitous drop in OD between 100-200 min after UV irradiation, indicating cell lysis (**Fig 2**). Following the drop in culture OD, there was a resumption of cell growth indicated by the increase in OD. Cells that were missing both SPβ and PBSX [(Ph(0); SAM614] had no detectable drop in OD following UV irradiation (**Fig 2**), indicating that the temperate phages SPβ and PBSX were responsible for cell lysis following UV irradiation.

**Figure 2.**
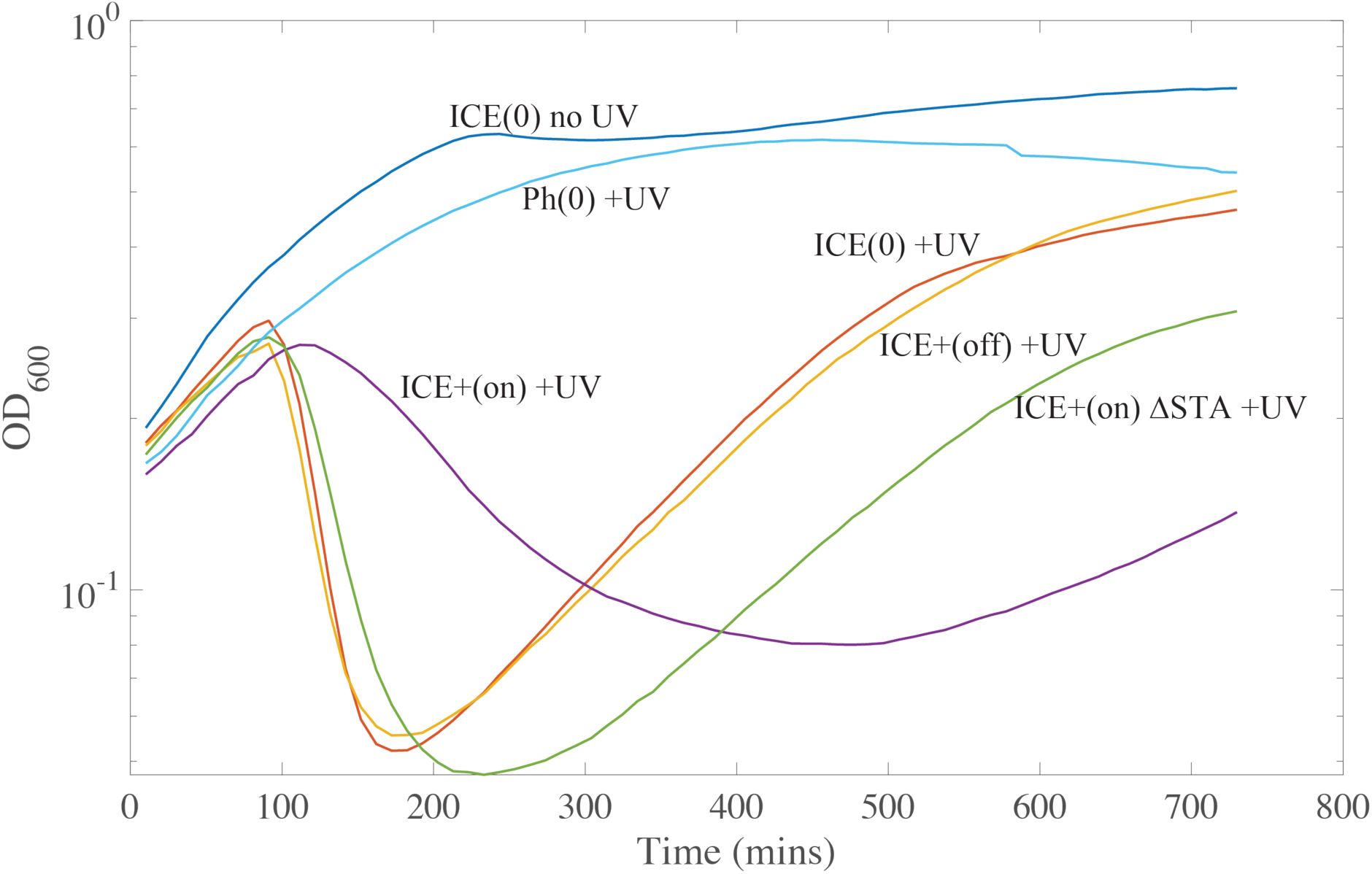
Expression of ICE*Bs1* prior to UV damage protects *B. subtilis* from phage-mediated lysis. Strains were grown at 37°C in minimal medium with 1% L-arabinose and grown to mid exponential phase (OD600 ∼0.2). Where indicated [ICE (on)], ICE*Bs1* was activated by inducing expression of *rapI* (from P*xyl*-*rapI*) with addition of 1% D-xylose 15 min prior to UV irradiation. Where indicated (+UV), cells were irradiated with UV light (Methods) and then grown shaking in a 96-well plate and growth was followed using a plate reader. OD600 is plotted versus time after irradiation. Strains containing ICE*Bs1* {ICE(+); SAM248} or cured of ICE*Bs1* [ICE(0); SAM591] were irradiated (+UV) or not (no UV). Ph(0) indicates a strain that is missing the two temperate phages SPβ and PBSX, but contains ICE*Bs1* (SAM614), activated for this experiment. ICE ΔSTA indicates cells that contain the ICE*Bs1* mutant that is missing *ramS, ramT,* and *ramA* (SAM601). Data from a representative experiment are shown. Similar results were obtained in at least three independent experiments.

Activation of ICE*Bs1* 15 min prior to UV irradiation (ICE+ (on); SAM248) caused a delay in cell lysis, a smaller drop in the OD of the culture, and a delay in the resumption of exponential growth (**Fig 2**). Based on these results, we conclude that activation of ICE*Bs1* conferred partial protection against phage-mediated lysis caused by the SOS response. As shown below, this is due to genes in ICE*Bs1* that modulate the host SOS response to DNA damage.

### A region of ICE*Bs1* that is necessary for ICE*Bs1-*mediated inhibition of DNA damage-induced cell lysis

Activation of ICE*Bs1* leads to robust expression of the polycistronic message produced from the promoter P*xis* (Auchtung *et al*., 2005; Auchtung *et al*., 2007), including expression of genes involved in excision, replication, and conjugation (**Fig 1A**). Because the observed inhibition of cell lysis required activation of ICE*Bs1*, the genes involved were likely expressed from P*xis* and not expressed when the element is integrated in the host chromosome and most element genes are repressed. We decided to initially focus on three genes with unknown function, *ramS*, *ramT*, and *ramA*, located downstream of *nicK* (encoding the element relaxase) (**Fig 1A**).

We deleted the *ramS, ramT, ramA* gene cluster (ΔSTA) in ICE*Bs1* and measured the effects of this mutant element on cell growth following UV irradiation, essentially as described above. In contrast to ICE*Bs1*+, we found that ICE*Bs1* ΔSTA (ICE+ ΔSTA; SAM601) did not confer protection against cell lysis and following UV irradiation (**Fig 2**). There was a precipitous drop in culture density, similar to or perhaps a bit more severe than that observed for cells without activation of ICE*Bs1* (**Fig 2**). The apparent increase in sensitivity to UV irradiation compared to cells without ICE*Bs1* or with an uninduced ICE*Bs1* might be due to production of ICE*Bs1* ssDNA during rolling-circle replication following induction of the element (Gigliani *et al*., 1993; Higashitani *et al*., 1995; Ruiz-Masó *et al*., 2015).

### Activation of ICE*Bs1* stimulates the host SOS response

We found that activation of ICE*Bs1* stimulated the host SOS response. We monitored the SOS response by measuring expression from the promoter of the SOS-inducible gene *yneA* fused to *lacZ* (P*yneA*-*lacZ*). *yneA* is repressed by LexA and de-repressed during the SOS response (Kawai *et al*., 2003; Goranov *et al*., 2006). We found that expression of P*yneA*-*lacZ* increased 120 minutes following activation of ICE*Bs1* (**Fig 3A-C**; ICE (on); AGH270). In contrast, expression was low in cells in which ICE*Bs1* had not been activated (**Fig 3A-C**; ICE (off); AGH270). These results indicate that activation of ICE*Bs1* caused activation of the host SOS response. This is likely due to production of ICE*Bs1* ssDNA during rolling-circle replication following induction of the element. Activation of ICE*Bs1* does not cause cell death (Menard and Grossman, 2013; Bean *et al*., 2022), indicating that stimulation of the SOS response by ICE*Bs1* is not sufficient to activate to activate the prophages SPβ and PBSX.

**Figure 3.**
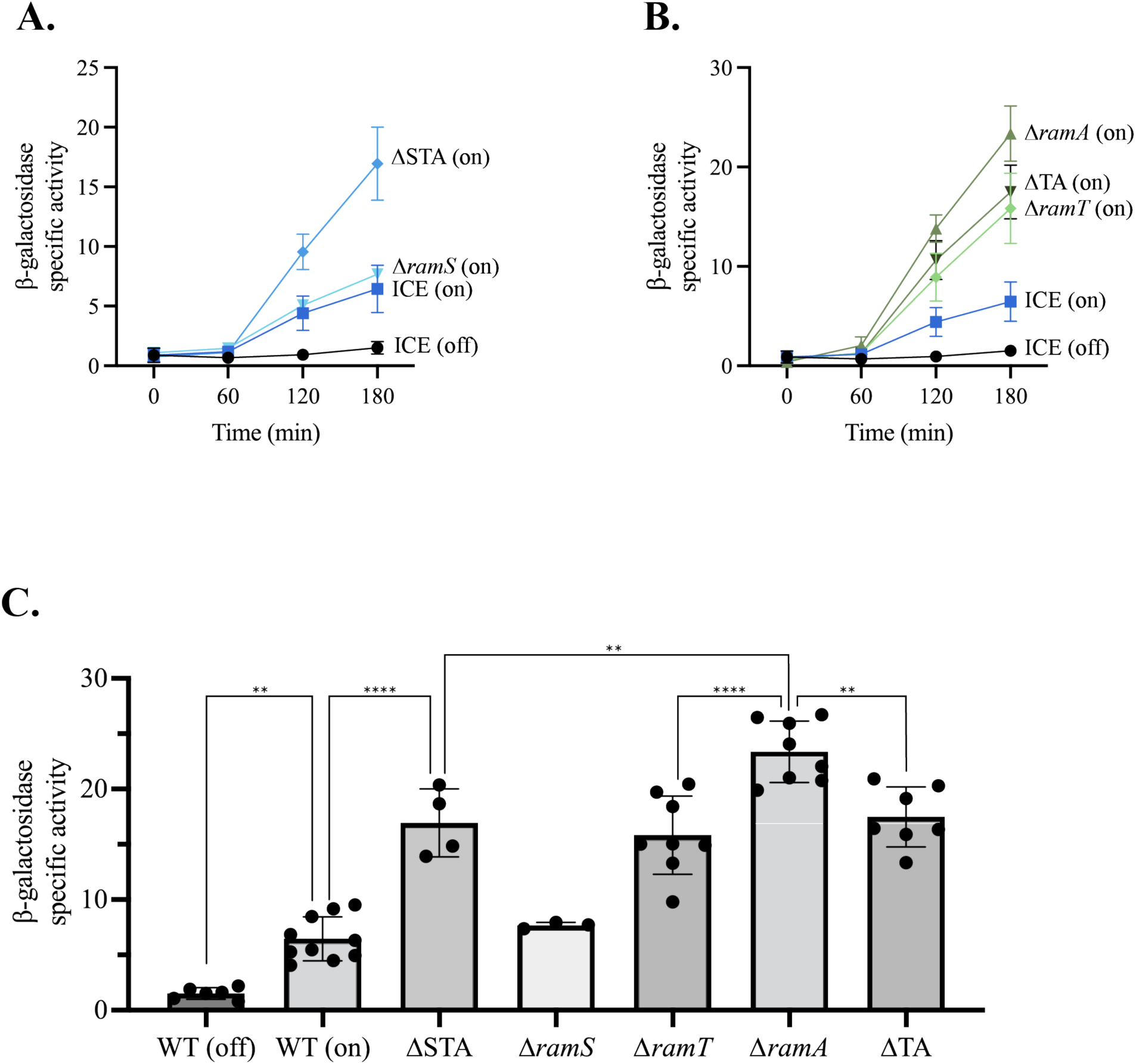
Effects of ICE*Bs1* and *ramSTA* on the host SOS response. Strains were grown at 37°C in minimal medium with 1% L-arabinose. ICE*Bs1* was activated where indicated, by inducing *rapI* (P*xyl*-*rapI*) with the addition 1% D-xylose when cells reached OD∼0.2. All strains have activated ICE*Bs1* except for ICE(off) samples. Samples were taken at indicated times and β-galactosidase specific activity was measured. All strains contain either wild-type ICE*Bs1* (ICE) or ICE*Bs1* with either single gene deletions (as indicated) or multiple gene deletions, as indicated: ΔTA (Δ*ramT-A*) and ΔSTA (Δ*ramS-T-A*). Data represent the mean of at least three independent biological replicates and error bars depict standard deviation. **A, B.** β-galactosidase specific activity vs time after activation of ICE*Bs1*. ICE (off) = AGH270 (black circles); ICE (on) = AGH270 (squares). **B.** Δ*ramS* = AGH241 (inverted triangles); ΔSTA = AGH239 (diamonds); Δ*ramT* = AGH207 and AGH353 (diamonds); ΔTA = AGH209 and AGH395 (inverted triangles); Δ*ramA* = AGH296 (triangles). **C.** β-galactosidase specific activity three hours after activation of ICE*Bs1*. Data are the same as presented in panels A and B. One-way ANOVA: P < 0.05 = *, P < 0.01 = **, P< 0.005 = ***, P < 0.001 = ****. Data that are not statistically different include: ICE (on) vs Δ*ramS*; Δ*STA* vs Δ*ramT* vs Δ*ramTA*.

### *ramT* and *ramA* function to inhibit the host SOS response

We found that deletion of *ramS, ramT,* and *ramA* (ΔSTA) from ICE*Bs1* caused an increase in the SOS response following activation of the element (**Fig 3A, C**; ΔSTA; AGH239). By three hours after activation, there was approximately twice as much β-galactosidase activity from P*yneA*-*lacZ* in cells with the ΔSTA mutation compared to that from the wild type (**Fig 3A, C**). These results indicate that the host SOS response was somehow reduced by one or more of the genes *ramS, ramT*, and *ramA*. Although RamS and RamT are 52.27% identical and 72.73% similar in a pairwise sequence alignment, we found that their functions are not redundant (see below).

#### ramS

Loss of *ramS* had little to no effect on activation of the SOS response (**Fig 3A, C**). That is, expression of P*yneA*-*lacZ* was virtually indistinguishable in strains with ICE*Bs1* wild type (AGH270) or ICE*Bs1* Δ*ramS* (AGH241) following activation of the element. Based on these results, we focused on *ramT* and *ramA*.

#### ramT and ramA

We found that deletion of either *ramT* (AGH207/AGH353) or *ramA* (AGH296) caused an increase in expression of P*yneA*-*lacZ* relative to that caused by activation of wild-type ICE*Bs1* (**Fig 3B, C**). Expression in the *ramA* mutant was consistently greater than that in the *ramT* mutant (**Fig 3B, C**). These results establish the genetic formalism that both RamA and RamT act is inhibitors of the host SOS response.

Expression of P*yneA*-*lacZ* in a *ramT ramA* (ΔTA, AGH209/AGH395) double mutant was less than that in the *ramA* single mutant (**Fig 3B, C**). This reduced expression caused by loss of *ramT* indicates that RamT is an activator (in a formal sense, irrespective of mechanism) of the host SOS response. In other words, the elevated SOS response in the *ramA* single mutant relative to the double mutant was due to the activating effect of *ramT*. Further, the phenotype of the *ramT ramA* double mutant was similar to that of the *ramT* single mutant (**Fig 3B, C**). That is, there was no discernable effect of *ramA* on the SOS response in the absence of *ramT*. This indicates that RamT likely functions in some way to stimulate or enable RamA to inhibit the SOS response and that RamA has no effect on the SOS response without RamT. This could be by activating or working together with RamA, or by affecting a cellular process that enables RamA to inhibit the SOS response.

To summarize the genetic interactions, the phenotypes of the *ramT* and *ramA* single and double mutants indicate that RamA functions to inhibit and RamT functions to both inhibit and activate the host SOS response, and that inhibition mediated by RamT is most likely through stimulating or enabling the repressive effects of RamA (**Fig 1B**).

### The SOS response is induced in a subpopulation of cells with activated ICE*Bs1*

The P*yneA*-*lacZ* reporter indicates gene expression on a population level and does not distinguish whether some or all of the cells were undergoing an SOS response. To determine the fraction of cells in a population that were undergoing SOS, we used a P*yneA*-*mNeongreen* reporter and fluorescence microscopy to monitor the SOS response in single cells.

We found that two hours after activation of ICE*Bs1,* approximately 18% of cells were fluorescent (**Fig. 4**; ICE+; SAM352), indicating that these cells had activated the SOS response. These results indicate that activation of ICE*Bs1* stimulated the SOS response in a subpopulation of cells.

**Figure 4.**
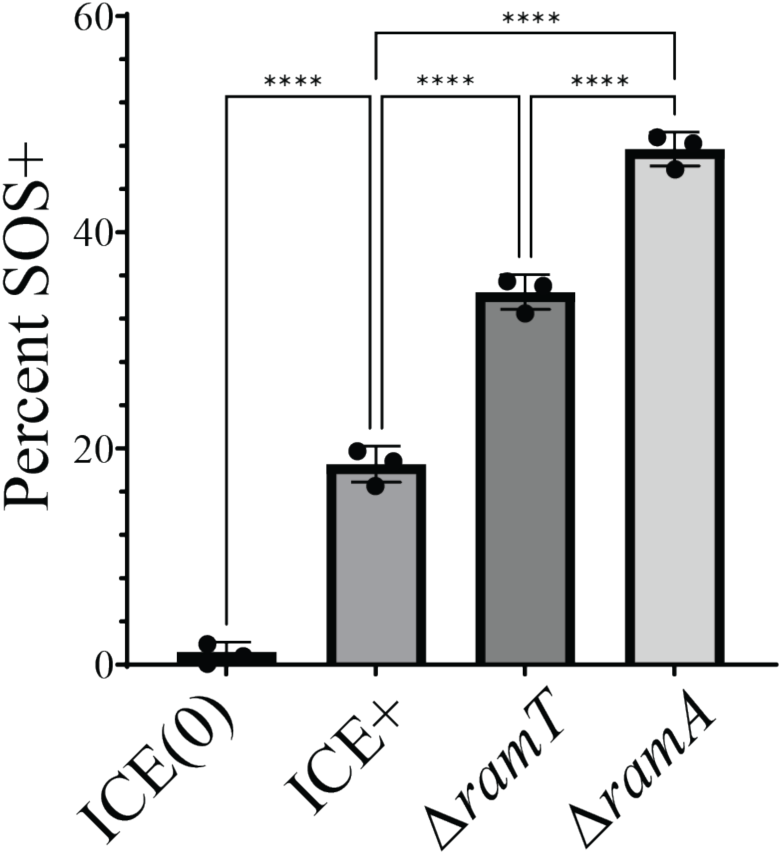
Activation of ICE*Bs1* induces the SOS response in a subpopulation of cells. Cells were grown at 37°C in minimal medium with 1% L-arabinose. ICE*Bs1* was activated by inducing *rapI* (P*xyl*-*rapI*) with the addition 1% D-xylose when cells reached OD600 of ∼0.2. All strains contained P*yneA*-*mNeongreen* as an indicator of the SOS response. The fraction of the population expressing PyneA-mNeongreen two hours after activation of ICE*Bs1* is shown. Data represent the mean of three independent biological replicates and error bars depict standard deviation. ICE(0) = SAM362; ICE+ = SAM352; Δ*ramT* = SAM955; Δ*ramA* = SAM957. Statistical significance: One-way ANOVA: P < 0.05 = *, P < 0.01 = **, P< 0.005 = ***, P < 0.001 = ****.

We found that *ramT* and *ramA* were involved in limiting the fraction of cells undergoing an SOS response following activation of ICE*Bs1*. Two hours after activation of ICE*Bs1* Δ*ramT*, approximately 35% of cells were expressing P*yneA*-*mNeongreen* (**Fig 4**; Δ*ramT*; SAM955), about 2-fold greater than that for wild-type ICE*Bs1*. Loss of *ramA* had a somewhat larger effect. Approximately 45-50% of cells were expressing P*yneA*-*mNeongreen* two hours after activation of ICE*Bs1* Δ*ramA* (**Fig 4**; Δ*ramA*; SAM957) about 2.6-fold greater than that for wild-type ICE*Bs1* (**Fig 4**; ICE+; SAM352), similar to the effects measured in the bulk population. (**Fig. 3 B, C**).

Together, our results indicate that activation of ICE*Bs1* stimulates the SOS response in a subpopulation of cells, and that *ramT* and *ramA* normally function to limit this activation. This inhibition of the SOS response caused by *ramT* and *ramA* is consistent with their roles in inhibiting host killing by the prophages SPβ and PBSX following UV irradiation.

### Formation of RecA filaments is inhibited by *ramT* and *ramA* during the SOS response

Binding of RecA to ssDNA following DNA damage leads to formation of RecA filaments along the ssDNA. We measured the levels and distribution of RecA-GFP in individual cells following treatment with mitomycin-C to induce the SOS response (**Fig 5A-C**) and summarized these data in demographs (**Fig 5D-H**). We used strains that were either missing ICE*Bs1* [ICE(0)] or cells that had a mini-ICE, containing all of the genes from *attL-ramA* (**Fig. 1A**), but missing the genes downstream of *ramA*, including those encoding the mating machinery. This eliminated the ability of the element to transfer and allowed us to focus on our genes of interest (*ramS-A*) and those involved in element replication, which generates ssDNA. Cells were sorted by length, with small cells at the top of the demographs and longer cells that are nearing or undergoing cell division at the bottom. The colors indicate the percentage of cells with RecA-GFP signal (red for higher percentage of cells, blue for lower percentage of cells) at the corresponding distance from the mid-cell (x-axis) The cumulative number of cells analyzed for each condition are indicated on y-axis and the distance from mid-cell on the x-axis.

**Figure 5.**
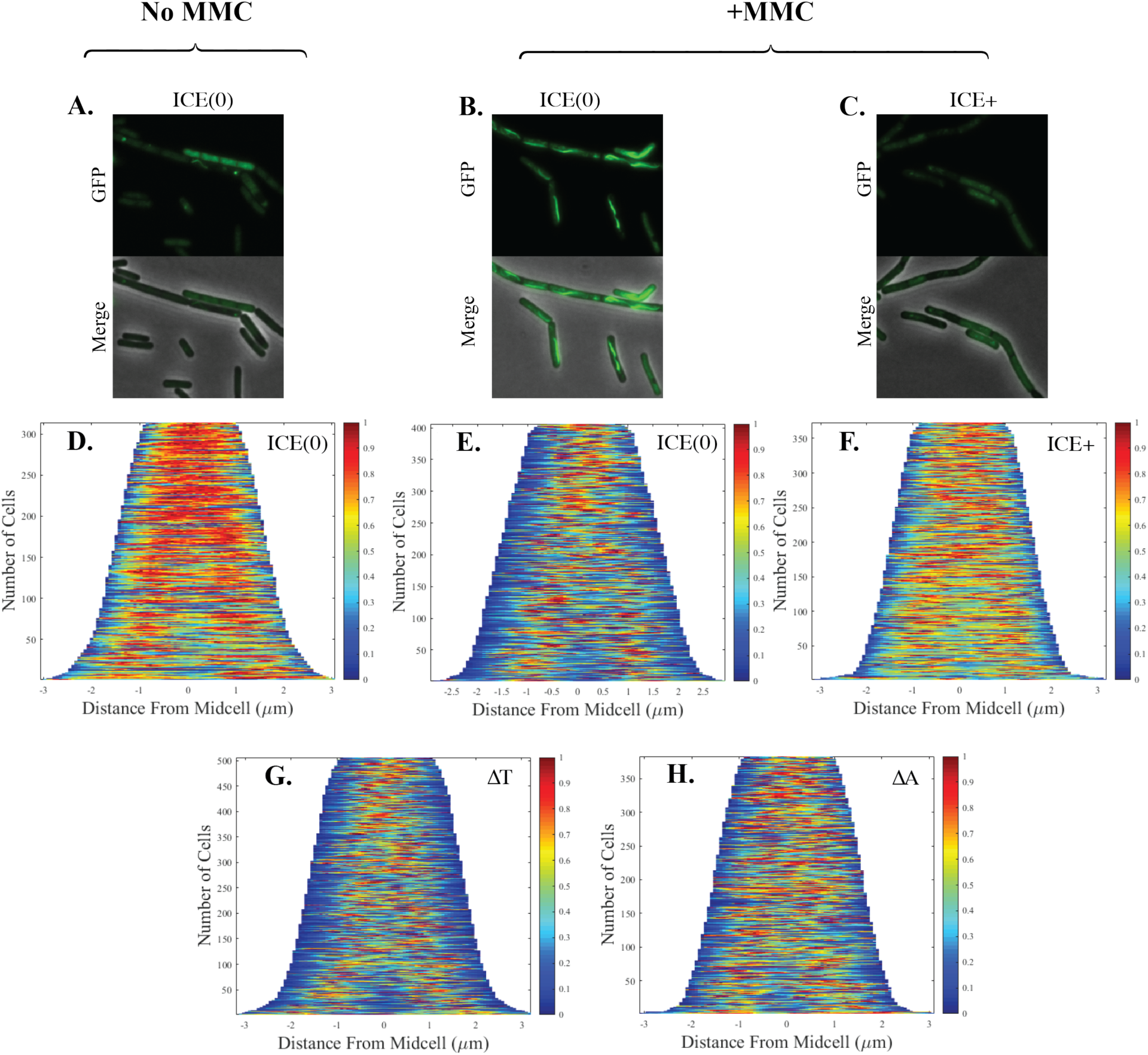
ICE*Bs1*-mediated inhibition of formation of RecA-GFP filaments. Strains were grown at 37°C in minimal medium with 1% L-arabinose to OD600 of ∼0.2, treated as indicate below, and samples taken for microscopy to visualize RecA-GFP. Exponentially growing cells without **(no MMC; A, D)** and 15 min after treatment with MMC **(+MMC; B, C, E-H)**. ICE*Bs1* was activated **(C, F-H)** by inducing *rapI* (P*xyl*-*rapI*) with the addition 1% D-xylose for 15 mins before the addition of MMC. ICE(0) indicates strains that were cured of ICE*Bs1*. ICE+, ΔT, and ΔA indicate strains containing mini-ICE elements (deleted for all the genes downstream of *ramA*), which contain are otherwise either wild type, Δ*ramT*, or Δ*ramA* respectively. At least three biological replicates, with six frames for each condition were analyzed. Demographs are from 30 mins after the addition of MMC. **A,B,C)** Representative micrographs of RecA-GFP filament formation under different conditions. **D-H)** Representative demographs of RecA-GFP filament formation. The fraction of cells containing RecA-GFP is indicated by the color scale (right y-axis) and the localization within the cell (determined as distance from midcell) is represented on the x-axis. The cumulative number of cells analyzed is indicated on left y-axis. **A-B,D-E)** ICE*Bs1*(0) = SAM591; **C,F)** WT ICE*Bs1* = SAM734; **G)** Δ*ramT* = SAM1003; **H)** Δ*ramA* = SAM1004.

In cells growing exponentially without ICE*Bs1* (and no mitomycin-C), RecA-GFP was distributed throughout the cell and did not form filaments or foci (**Fig 5A**; ICE(0), no MMC; SAM591) similar to what has been described previously (Simmons *et al*., 2007). In the demograph, this is represented by a general red signal across much of the length of the cell, typically centered around mid-cell in smaller cells and the cell quarters (which will become mid-cell following division) in larger cells (**Fig 5D**; ICE(0), no MMC; SAM591).

Treatment of cultures with mitomycin C for 30 min resulted in the formation of RecA-GFP filaments in strains without ICE*Bs1* (**Fig. 5B**; ICE(0), +MMC; SAM591). This is seen as the loss of much of the red throughout the cells, and a more focused signal near mid-cell (or the cell quarters), and a concomitant increase in blue in the demographs (**Fig. 5E**; ICE(0), +MMC; SAM591). The mid-cell position is likely indicative of the location of the replisome and consistent with previous findings (Simmons *et al*., 2007).

Strikingly, formation of RecA-GFP filaments was quite different in strains in which ICE*Bs1* had been activated for 15 min before treatment with mitomycin-C (**Fig. 5C, F**; ICE+, +MMC; SAM734). The distribution of RecA-GFP was between that seen in ICE-cured cells without (**Fig 3D**) and with mitomycin C (**Fig 3E**). These results indicate that expression of ICE*Bs1* was inhibiting the formation of RecA-GFP filaments.

We found that both *ramT* and *ramA* were required for the ICE*Bs1*-mediated inhibition of RecA-GFP filament formation. The pattern of RecA-GFP filaments in a population of cells in which ICE*Bs1* Δ*ramT* (**Fig 5G**; ΔT, +MMC; SAM1003) or ICE*Bs1* Δ*ramA* (**Fig 5H**; ΔA, +MMC; SAM1004) had been activated for 15 min prior to MMC treatment was largely similar to that in cells without activation of ICE*Bs1* (**Fig 5E**).

Together, our results indicate that *ramT* and *ramA* inhibit the cellular SOS response, and that this inhibition is mediated, at least in part, by either directly or indirectly affecting formation of RecA filaments.

## Discussion

Our work demonstrates that ICE*Bs1* modulates the host DNA damage (SOS) response by inhibiting accumulation of RecA filaments, either directly or indirectly. We found that activation of ICE*Bs1*, without external damaging agents, was sufficient to activate the SOS response in a subpopulation of cells and that the ICE*Bs1* products RamT and RamA attenuate this response by reducing the presence of RecA filaments. Further, activation of ICE*Bs1* prior to treatment with a DNA damaging agent also reduced the SOS response, reducing activation of resident temperate phages, thereby limiting phage-mediated cell death. This effect was also mediated by *ramT* and *ramA*. Our results show that RamA inhibits and RamT both inhibits and activates the host SOS response, and the inhibition mediated by RamT is likely achieved by stimulating inhibition by RamA (**Fig 1B**).

Our experiments were done with a population of ‘donor’ cells, all of which contained the element. We suspect that the inhibition of SOS also occurs in transconjugants shortly after transfer of ICE*Bs1* to a new host. The DNA that is transferred by type IV secretion systems is linear ssDNA, a substrate for Ssb and indicator of DNA damage. After entry into a new host and the conversion of the linear ssDNA to circular dsDNA, *ramT* and *ramA* will be expressed from the strong element promoter P*xis*. We postulate that this expression will help limit the SOS response in transconjugants, before P*xis* is repressed and the element integrates into the chromosome. Additionally, *ramT* and *ramA* are downstream from the single strand origin of replication (*sso*) in ICE*Bs1*. The ICE *sso* is likely recognized by the host RNA polymerase to make the primer for DNA synthesis (Wright *et al*., 2015). Since *ramT* and *ramA* are downstream from the *sso*, we suspect that these two genes might be transcribed shortly after entry of the ICE ssDNA.

We hypothesize that the ability of ICE*Bs1* to inhibit activation of the SOS response allows the element to remain activated with the potential to transfer to other cells for extended periods of time, while reducing fitness costs to its host. Active rolling-circle replication produces a considerable amount of ssDNA bound by Ssb (Gigliani *et al*., 1993; Higashitani *et al*., 1995; Ruiz-Masó *et al*., 2015). This is typically a signal of DNA damage and causes activation of the SOS response and subsequent inhibition of cell division. Limiting the population of cells undergoing the SOS response likely allows many cells to continue to divide while the element is extrachromosomal and in multicopy. We suspect that inhibition of the SOS response is a conserved property of other ICEs that undergo robust rolling-circle replication.

Many of the *Bacillus* strains that contain ICE*Bs1* or ICE*Bs1*-like elements also harbor prophages (both functional and defective) that are activated by the SOS response, as is ICE*Bs1*. In our lab strains, the SOS response elicited by activation of ICE*Bs1* is not sufficient to activate the resident prophages SPβ and PBSX, even in the absence of *ramT* and *ramA*. However, activation of ICE*Bs1* prior to more robust DNA damage reduces activation of the resident prophages. If an element is activated at a lower threshold of DNA damage than the resident prophages, then this could help protect cells from phage-mediated killing. Further, there are likely other temperate phages that are more sensitive to DNA damage signals than SPβ or PBSX. We suspect that ICE-mediated inhibition of SOS likely protects cells from activation of those phages.

### Multiple ways ICE*Bs1* affects the physiology of host cells

Inhibition of RecA filaments and the SOS response is one of at least four ways in which ICE*Bs1* affects host cell physiology and/or fitness. The three other known ways include: 1) The ICE*Bs1* gene *devI* causes a delay in sporulation and biofilm formation when the element is active (excised from the chromosome with its genes expressed) and allows extra cell divisions to ICE*Bs1*-containing cells. This delay in development enables those cells to out-compete neighboring cells without the element (Jones *et al*., 2021). 2) ICE*Bs1* has an abortive infection mechanism mediated by *spbK* that protects ICE*Bs1* host populations from predation by the phage SPβ (Johnson *et al*., 2022). *spbK* is expressed independently of other ICE*Bs1* genes, even when the element is integrated in the host chromosome. 3) The product of the ICE*Bs1* gene *yddJ* inhibits the activity of the cognate conjugation machinery in other cells. This ‘exclusion’ mechanism limits excessive conjugative transfer and its potentially lethal effects on the cell envelope (Avello *et al*., 2019). Together these strategies play an important role in the transfer and maintenance of ICE*Bs1* and increases the competitive advantage of both the element and its host.

### Other modulators of the SOS response encoded by mobile genetic elements

ICE*Bs1* is not the only mobile genetic element that has genes for inhibiting the host SOS response. One of the best understood element-encoded inhibitors of the host SOS response is PsiB, encoded by the F plasmid of *E. coli* and other related conjugative plasmids. *psiB* is expressed in transconjugants and its product binds to and inhibits RecA, thereby limiting the SOS response and activation of resident prophages (Bagdasarian *et al*., 1986; Bagdasarian *et al*., 1992; Althorpe *et al*., 1999; Petrova *et al*., 2009; Baharoglu *et al*., 2010).

RamT and RamA, while inhibiting RecA filaments, are not similar in sequence to PsiB, or any other characterized protein. Further, PsiB appears to act only in transconjugants (Jones *et al*., 1992; Althorpe *et al*., 1999), whereas RamT and RamA act in donors, and we suspect also act in transconjugants. In these ways, the mechanisms by which RamT and RamA function are likely to be quite different from that of PsiB.

Some bacteriophages also have mechanisms to modulate the host SOS response and recombination systems. For example, bacteriophage lambda encodes the Redβ recombination system (*exo*: exonuclease, *bet*: strand-annealing protein, *gam*: RecBCD nuclease inhibitor) along with *orf* (recombination mediator), and *rap* (Holliday junction endonuclease). Together these proteins are sufficient to perform many of the host functions of DNA damage repair, including homologous recombination, recognition of SSB-bound ssDNA, and Holliday junction resolution (Poteete, 2001; Poteete, 2004; Casjens and Hendrix, 2015). Though these genes, in large part, are there to complete a specific step in the lambda phage life cycle, some inhibit activation of the SOS response. Gam inhibits the activity of the exonuclease RecBCD (essential for double-strand break repair) when lambda switches from theta to rolling-circle replication (Zissler *et al*., 1971; Smith, 2012), a crucial point when lambda is most likely to activate the SOS response.

Some phages act directly on LexA to inhibit the host SOS response. For example, the tectiviral temperate phage GIL01 of *B. thuringiensis* encodes a small protein (gp7) that forms a stable complex with LexA, enhancing binding of LexA to SOS boxes and inhibiting RecA-mediated autocleavage required to activate the SOS response (Fornelos *et al*., 2015). GIL01 dampens the host SOS response and activation of other co-resident prophages to ensure that it is able to become active and replicate before the host activates the SOS response and derepresses other phages (Fornelos *et al*., 2016; Brady *et al*., 2021; Pavlin *et al*., 2022).

Determining how ICEs modulate the physiology of their hosts is critical for understanding the complex relationships between different mobile elements within a given host. These interactions are continually evolving and conjugative elements have multiple ways to help protect their bacterial hosts from predation by phages, including modulating the SOS response.

## Methods and Materials

### Media and growth conditions

Cells were grown in LB medium (usually for strain construction) or S750 defined minimal medium (Jaacks *et al*., 1989). For experiments, all strains were colony purified from frozen (−80°C) glycerol stocks on LB agar plates with the appropriate antibiotics and cells from a single colony were inoculated into S750 minimal medium with 1% L-arabinose w/v as the carbon source and required amino acids (40 μg/ml phenylalanine and 40 μg/ml tryptophan). Cells were grown to mid-exponential phase, then diluted to an OD600 of 0.025 and grown at 37°C with shaking until an OD600 of ∼0.2. ICE*Bs1* was activated by adding 1% D-xylose to induce expression of Pxyl-*rapI*. RapI causes the ICE*Bs1*-encoded protease ImmA to cleave the repressor ImmR, thereby derepressing ICE*Bs1* gene expression (**Fig 1A**) (Auchtung *et al*., 2007; Bose *et al*., 2008; Bose and Grossman, 2011). Antibiotics used at the following concentrations: 5 μg/ml kanamycin, 10 μg/ml tetracycline, 100 μg/ml spectinomycin, 5 μg/ml chloramphenicol, and a combination of erythromycin at 0.5 µg/ml and lincomycin at 12.5 µg/ml to select for macrolide-lincosamide-streptogramin (*mls*) resistance.

### Bacterial strains and alleles

Strains were constructed by natural transformation. All strains (**Table 1**) are derivatives of JH642 (AG174; (Smith *et al*., 2014) and contain *trpC* and *pheA* mutations (not indicated). Strains cured of ICE*Bs1* are indicated as ICE*Bs1*^0^. Strains containing either *amyE*::[P*xyl*-r*apI*), *spc*] (Berkmen *et al*., 2010) or *lacA*::[P*xyl*-*rapI*), *tet*] (Lee *et al*., 2010) were used to activate expression of ICE*Bs1* (following addition of xylose). Most ICE*Bs1* strains contained a kanamycin-resistance gene [Δ(*rapI-phrI*)*342*::*kan*] (Auchtung *et al*., 2005) or a derivative, to allow selection for the element. The *recA*:{(*recA-gfp*) *spc*} (Simmons *et al*., 2007), the Ph(0) ΔPBSX::lox ΔSPβ::lox (Schons-Fonseca *et al*., 2022), the *PyneA-mNeongreen* (McKeithen-Mead and Grossman, 2023), and the *PyneA-lacZ* (Biller *et al*., 2011); obtained from the Bacillus Genetic Stock Center) alleles have all previously been described previously.

**Table 1.**
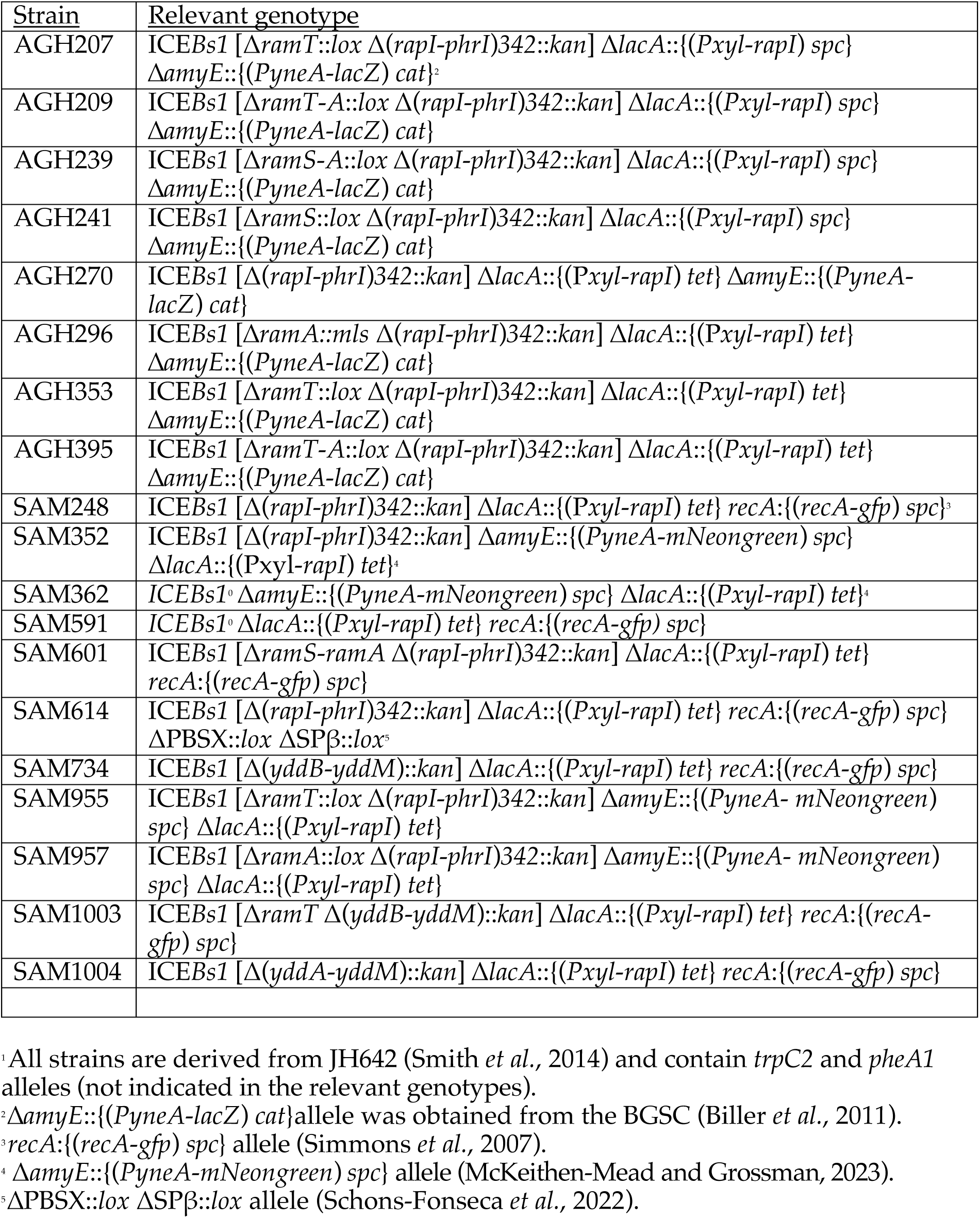
*B. subtilis* strains used^1^.

#### Δ(ramS-ramA)

The three genes *ramS, ramT*, and *ramA* were deleted in two steps, first by integrating a plasmid and then by screening for loss of the integrated plasmid and the presence of the deletion allele. A fragment from ∼500 bp upstream of *ramS* to the first 12 bp of *ramS,* and a second fragment from the last 42 bp of *ramA* (to maintain the start of *conB*, which overlaps the end of *ramA*) to ∼500 bp downstream of *ramA* were amplified by PCR and assembled into pCAL1422 (Thomas *et al*., 2013; a plasmid that contains *E. coli lacZ*) by isothermal assembly (Gibson *et al*., 2009). The resulting plasmid pELC1582, was integrated into the chromosome by single-crossover recombination. Transformants were screened for loss of *lacZ*, indicating loss of the integrated plasmid, and PCR was used to identify a clone containing the Δ(*ramS ramT ramA*) allele (indicated as Δ*ramS-ramA* or ΔSTA).

#### ΔramS::lox, ΔramT::lox, ΔramA::lox, and ΔramT-ramA::lox

Deletions of individual genes and the *ramT-ramA* double deletion with *lox* insertions were made by introducing an antibiotic resistance gene (*cat*) flanked by *lox* sites into each gene followed by Cre-mediated recombination to remove the antibiotic resistance gene and leaving a *lox* insertion in place of the coding sequence. Regions from about ∼1 kb upstream to ∼3 bp in each gene and from the stop codon to ∼1 kb downstream of each gene, except for *ramA*, were amplified by PCR. For *ramA*, 28 bp the 3’-end of the gene were included to maintain the *conB* open reading frame. For the *ramT-ramA* double deletion, a region from ∼1 kb upstream of *ramT* to 4 bp into *ramT* and a region from the last 40 bp at the 3’-end of *ramA* to ∼1 kb downstream of *ramA* were used. The *cat* cassette from pGemcat (Harwood and Cutting, 1990) was amplified with primers that added *lox71* and *lox66* sites. For each gene, the PCR fragments from upstream and downstream sequences were assembled on either side of *cat* (with the *lox* sites) using linear isothermal assembly (Gibson *et al*., 2009). The assembled fragment was integrated into the chromosome by double-crossover and selecting for resistance to chloramphenicol. *cat* was removed by Cre-mediated recombination between the *lox* sites following introducing (by transformation) of a temperature-sensitive plasmid expressing the Cre recombinase (pSAM097). Strains were cured of pSAM097 by shifting cells to a non-permissive temperature and allowing for growth without selection for the plasmid. Diagnostic PCR and sequencing were used to confirm each allele.

#### ΔramA::mls

The *ramA* deletion-insertion replaced most of the open reading frame of *ramA*, leaving intact the first 10 bp and last 40 bp of the open reading frame, with an antibiotic resistance cassette (*mls*) using linear isothermal assembly. Regions flanking ∼1 kb upstream and downstream of these boundaries and the *mls* cassette from pDG795 (Guérout-Fleury *et al*., 1996) were amplified by PCR, fused with linear isothermal assembly (Gibson *et al*., 2009), and used for transformation.

#### Mini-ICE strains for RecA-GFP localization

Mini-ICE elements were constructed by deleting *yddB-yddM,* either with or without *ramA* or *ramT*, and introducing a kanamycin resistance cassette. For the wild-type mini-ICE, regions from ∼1kb upstream of *yddB* to the stop codon of *ramA*, and from the last 5 bp of *yddM* to ∼1kb downstream of *yddM* were used. For the Δ*ram*A mini-ICE, regions from ∼1kb upstream of *ramA* to 24 bp downstream of *ramT* and from the last 5 bp of *yddM* to ∼1kb downstream of *yddM* were used. For the Δ*ramT* mini-ICE, regions from ∼1kb upstream of the start *ramT* to 12 bp after the end of *ramS*, from the end of *ramT* to the end of *ramA*, and from the last 5 bp of *yddM* to ∼1kb downstream of *yddM* were used. The kanamycin resistance cassette was obtained from pGK67 (Auchtung *et al*., 2005). These fragments were amplified using PCE and combined using linear isothermal assembly (Gibson *et al*., 2009), and transformed into JH642, selecting for kanamycin resistance. Additional alleles (described above) were added to generate the final strains.

### UV irradiation and effects on cell growth

Cells were grown to mid-exponential phase in defined minimal medium with 1% L-arabinose and grown to mid exponential phase (OD600 ∼0.2). D-xylose (1%) was added to activate ICE*Bs1* where indicated. After 15 mins, 600 µl of each culture was placed into a sterile plastic petri dish and samples were irradiated with UV light (5 J/m^2^) using a Stratalinker UV Crosslinker. Aliquots (180 µl) of each culture were then grown in a 96-well plate (96-Well Non-Treated Plates, GenClone) with three technical replicates for each culture. Cultures in the 96-well plates were grown at 37°C with continuous double orbital shaking at 807 cpm in a Biotek Synergy H1 plate reader and OD600 readings taken over the course of the experiment, typically for 10-12 hrs.

### β-galactosidase assays

Cells were grown at 37°C in defined minimal medium with 1% L-arabinose and grown to mid exponential phase (OD600 ∼0.2). D-xylose (1%) was added to activate ICE*Bs1* where indicated. Samples were taken at indicated timepoints. Cells were permeabilized with 15 µl of toluene and β-galactosidase specific activity was determined [(ΔA420 per min per ml of culture per OD^600^ unit) × 1000] essentially as described (Miller, 1972) after pelleting cell debris.

### Microscopy and image analysis

Imaging of single cells was done with an inverted microscope (Nikon, Ti-E) with a motorized stage and a CoolSnap HQ2 camera (Photometrics). A Nikon Intensilight mercury illuminator was used with appropriate sets of excitation and emission filters (filter set 49002 for GFP, Chroma). Cells were spotted on agarose pads set in a homemade incubation chamber made by stacking two sealable Frame-Seal Incubation Chambers (BIO-RAD). Agarose pads contained 1.5% Ultrapure agarose (Thermo Fisher) dissolved in Spizizen salts medium (Harwood and Cutting, 1990).

Images and videos were processed with Fiji ImageJ (Schindelin *et al*., 2012). All images for fluorescent channels were subjected to background subtraction using 10 or 20 pixels (10 pixels was used for images with RecA-GFP filamentation). Cell meshes were acquired using Oufti (Paintdakhi *et al*., 2016). Meshes were analyzed in MATLAB using custom scripts.

Cells expressing mNeongreen were gated by comparing fluorescence to that of a strain containing the reporter but that did not contain ICE*Bs1* [ICE*Bs1*(0)].

## Acknowledgments

We thank Emily Bean for the Δ*ramS-ramA* allele and the BGSC for the *PyneA-lacZ* allele. Research reported here was supported, in part, by grants from the National Institute of General Medical Sciences of the National Institutes of Health under award number R01 GM050895, R35 GM122538 and R35 GM148343 to ADG. SAM was supported, in part, by the NIGMS predoctoral training grant T32 GM007287 and a MathWorks Science Fellowship. The funders had no role in study design, data collection and analysis, decision to publish, or preparation of the manuscript.

## Notes

### Competing Interest Statement

The authors have declared no competing interest.

